# Effects of ancestry, agriculture, and lactase persistence on the stature of prehistoric Europeans

**DOI:** 10.1101/2025.07.11.664181

**Authors:** Samantha L Cox, Kaeli Kaymak-Loveless, Carson Shin, Timka Alihodžić, Kurt W. Alt, Nadezhda Atanassova, Dider Binder, Morana Čaušević-Bully, Alexander Chohadzhiev, Stefan Chokhadzhiev, Henri Duday, Bisserka Gaydarska, Anahit Khudaverdyan, Rafael Micó Perez, Nicole Nicklisch, Mario Novak, Camila Oliart Caravatti, Hélène Réveillas, Maïté Rivollat, Stephane Rottier, Domagoj Tončinić, Steve Zaüner, Iain Mathieson

## Abstract

Ancient DNA has revolutionized our understanding of human evolutionary history, but studies focusing solely on genetic variation tell an incomplete story by neglecting phenotypic outcomes. The relationships between genotype and phenotype can change over time, making it desirable to study them directly in ancient populations rather than present-day data. Here, we present a large-scale integration of ancient genomic and phenotypic data, analyzing femur length as a proxy for stature in 568 individuals with published whole-genome ancient DNA data across western Eurasia. Polygenic scores derived from modern European and East Asian genome-wide association studies retain predictive power in ancient populations, explaining up to 10% of phenotypic variance. Contrary to longstanding archaeological hypotheses, we find that Neolithic populations were only modestly shorter than preceding Mesolithic groups, with differences at least partly attributable to genetic rather than environmental factors, challenging narratives of systematic stature decline following the transition to agriculture. Finally, we find that the lactase persistence allele had a large positive effect on stature in ancient individuals (0.24 standard deviations), even though it shows no association with height in modern populations. This gene-environment interaction highlights the limitation of using present-day genetic data to infer past phenotypic relationships. Our results underscore the value of integrating genetic and morphological data from ancient populations to reconstruct the dynamics of human adaptation.

## 1 Introduction

Adaptation and evolution reflect the cumulative outcomes of gene-environment interactions that shape phenotypic development within lifetimes and across generations. The genetics of these fundamental processes can be difficult to study in human populations mainly because present-day populations only retain the scars of past events, giving little insight into the process itself. Revealing the evolutionary history of complex traits necessitates the use of ancient DNA; however, there is less ancient data available and few quantifiable phenotypes with which genetic results can be verified.

Human stature represents an ideal model system for bridging this gap. As one of the most heritable complex traits, stature is both easily measured in skeletal remains and extensively characterized in modern genome-wide association studies. Its high polygenicity and environmental sensitivity make it particularly valuable for understanding how complex traits respond to changing selective pressures and how gene-environment interactions shift over time. Moreover, stature serves as a key indicator of population health and nutrition, making it central to debates about the major transitions of human prehistory.

One of the major environmental shifts in human prehistory was the global transition from hunting and gathering to farming, the causes and effects which have been a topic of intense study and debate for decades. One long standing hypothesis, first formalized by Armelagos and Cohen [1], posits that farming provided a stable environment that fueled population growth, but came at the cost of decreased individual health and a lower standard of living. Consequently, numerous researchers have investigated changes in European health between the Mesolithic and Neolithic, often using stature as a proxy, with conflicting results [e.g., 2, 3, 4, 5, 6, 7, 8]. Further, studies that do find stature decreases associated with the advent of farming have generally not accounted for the dramatic shifts in genetic ancestry that coincide with this major cultural and technological transition [9, 10] (with exceptions [11]). While anthropologists have always recognized that human biology is the result of complex interactions between genetics, culture, and environment, separating these effects has proved challenging.

Ancient DNA provides one potential approach. Several studies have combined GWAS data with ancient DNA to compute polygenic scores and track the evolution of stature in ancient individuals across time [12, 13], and to compare with changes in skeletal stature [14]. However, this approach has two major limitations. First, polygenic scores do not transfer well among present-day ancestry groups [15] and thus, presumably, not between present-day and ancient populations. Second, we do not know the relevant environmental effects and cannot account for their change through time. Though we assume a change in polygenic score reflects a change in phenotype, it can equally signal a shift in environment with no change in the resulting phenotype [16]. On the other hand, environmental perturbations can also lead to large changes in height without genetic change in both ancient [17] and present-day populations [18]. Finally, since questions about the magnitude and prevalence of gene-environment interactions in present-day populations remain largely unanswered, it is still unknown whether the relationships between genes, environment and phenotype are the same in ancient populations.

Here, we investigate the evolutionary history of human height using maximum femur length as a proxy for attained stature. We collected a dataset consisting of maximum femur length and ancient DNA from 568 individuals from Eurasia, dated between 38,000 and 600 years BP. We then leveraged this combination of data to infer the contribution of environment to height across Neolithic agricultural communities and later societies, measure individual SNP effects on height in ancient individuals, and investigate the interaction between lactase persistence and environment. Our study represents the first systematic analysis of this scope in an ancient population, resulting in the first empirical evidence of gene-environment interaction in an ancient human sample.

## 2 Results

We collected femur lengths for 568 individuals with previously published ancient DNA (Supplementary Table 1). The bulk of our data are concentrated in Western Eurasia and dated after 8000 BP (n=513/568) (Fig. 1A). We either measured femur lengths directly or reverse engineered them from published stature estimations in order to remove error associated with combining stature estimates using different estimation methods. We previously showed that genotype imputation can improve PRS performance in low-coverage ancient samples [19]. Given the low coverage of our sample (median=0.515×), we imputed diploid genotypes using a previously published aDNA imputation pipeline [20, 21]. This causes many of our samples to shift consistently in PC space (Fig. 1B).

**Figure 1:**
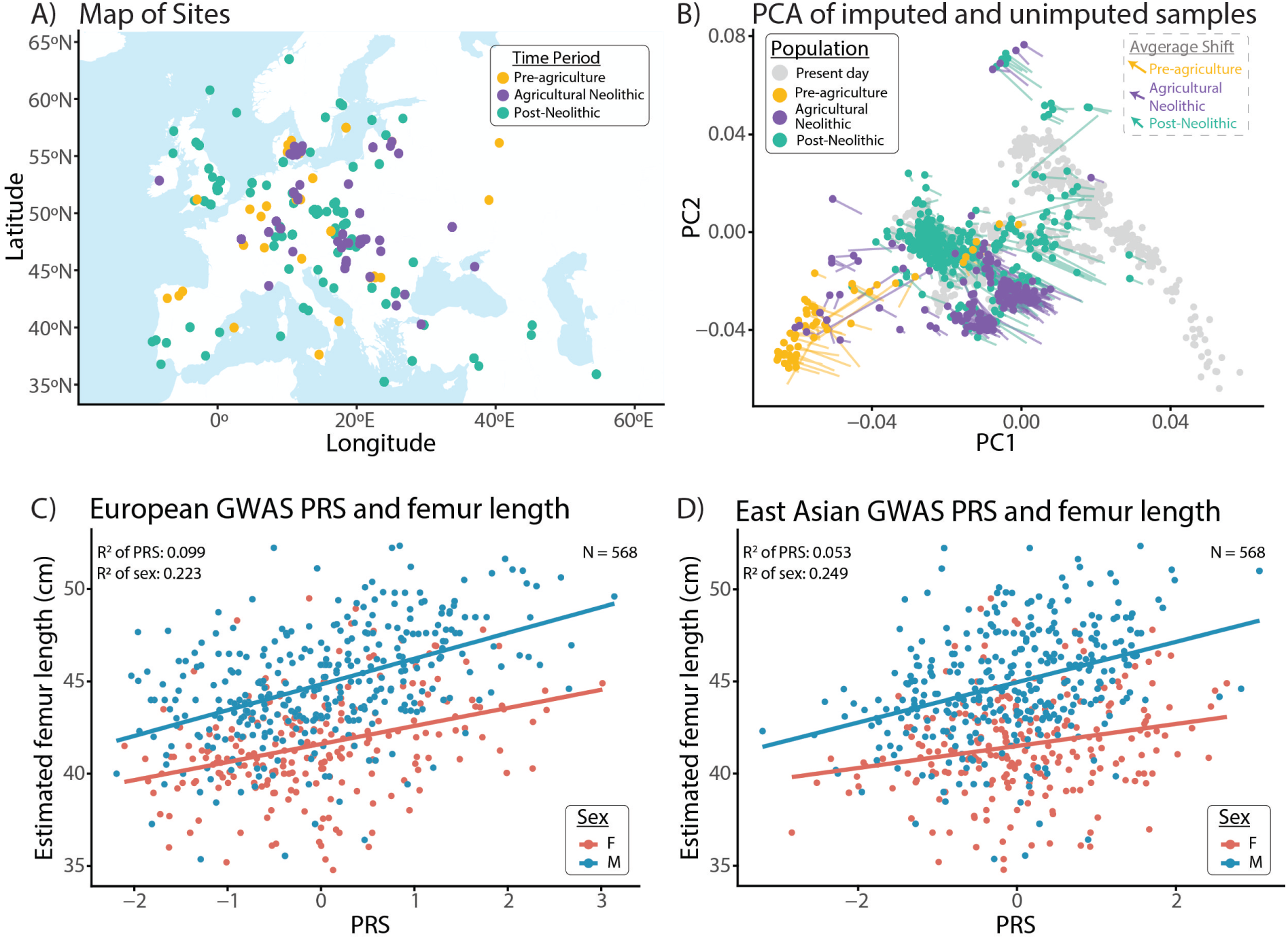
Sample characteristics. A) Distribution of samples across Europe. Populations were defined based on the published archaeological evaluation of each site (see Methods); B) PCA showing the relationship between imputed (colored points) and unimputed samples projected into present-day European (Human Origins) PC space (gray points). Tails indicate the original location of the unimputed point for each individual. Dashed box in the top right shows the average shift distance and direction in PC space for samples in each population, arrows are shown to scale; C) relationship between PRS and femur length by sex for the best performing European PRS calculated using 1240k imputed genomes and [22] European GWAS summary statistics; D) relationship between PRS and femur length by sex for the best performing unconfounded PRS calculated using the HapMap3 imputed genomes and [22] East Asian meta-GWAS summary statistics.

### 2.1 Identifying the optimal PRS for individual and population comparisons

We tested PRS constructed using clumping and thresholding on 1240k SNPs from several different source GWAS on both pseudohaploid and imputed genotype data: UK Biobank [23], East Asian [24], multi-ancestry [22] meta-analyses and between-sibling GWAS [25], as well as pre-trained PRS weights on HapMap3 SNPs [22]. To evaluate performance, we used linear regression of femur length on sex and four principal components, sample age and age^2^, calculating the incremental *R*^2^ when we add the PRS to the covariates. As our sample is largely from Europe, we were concerned that spurious associations between genetics and environment, known as population stratification, in the GWAS could confound results from PRS trained on present-day European ancestry populations [26, 27]. For this reason, we wanted to select two PRS for downstream analysis. The first, using European-derived summary statistics that maximized the incremental *R*^2^, but could be confounded by stratification, and the second from an independent population that should be unconfounded by European population structure but has lower predictive power.

We found that the overall optimal PRS used European (EUR) summary statistics [22] and clumping and thresholding on imputed 1240k SNPs (*R*^2^ = 0.099; Fig. 1C). This is about four-fold lower than the *R*^2^ obtained in present-day European individuals [22], likely due to lower data quality or increased distance to the training population from the ancient samples [28]. The optimal unconfounded approach was to use pre-trained East Asian (EAS) PRS weights [22] with imputed genotyped data on HapMap3 (HM3) SNPs (*R*^2^ = 0.053; Fig. 1D). For subsequent analyses we therefore use the Yengo EUR 1240K PRS when we want to maximize individual prediction, and the Yengo EAS HM3 PRS when we want to ensure our results are robust to population stratification, for example to compare population means.

**Table 1:** Comparison of R^2^ values with different PRS and imputation methods; *GWAS unconfounded by European population statification; red: highest European PRS R^2^; blue: highest unconfounded PRS R^2^.

| | PRS | pseudohaploid $R^2$ | imputed $R^2$ |
| --- | --- | --- | --- |
| 1240k SNPS<br>C&T | UK Biobank (EUR) | 0.065 | 0.088 |
|  | Chen et al., 2023 (EAS)* | 0.030 | 0.027 |
|  | Tan et al., 2024 (Sibs)* | 0.003 | 0.026 |
|  | Yengo et al., 2022 (ALL) | 0.066 | 0.097 |
|  | Yengo et al., 2022 (EUR) | 0.064 | 0.099 |
|  | Yengo et al., 2022 (EAS)* | 0.015 | 0.018 |
| HM3 SNPS<br>Yengo PGS | Yengo et al., 2022 (ALL) | - | 0.057 |
|  | Yengo et al., 2022 (EUR) | - | 0.084 |
|  | Yengo et al., 2022 (EAS)* | - | 0.053 |

### 2.2 The Neolithic transition had limited effect on stature

A long-standing hypothesis states that Neolithic populations were shorter than both the preceding and subsequent populations due to the negative health consequences of the transition to agriculture [1, 29]. This suggests that higher rates of population growth in early agricultural populations came at the cost of impaired individual health. However, phenotypic changes cannot necessarily be attributed to agriculture since the Neolithic transition in Europe coincides with major transitions in ancestry which could also contribute to changes in stature [30, 10, 14]. In principle, this can be resolved by comparing genetically predicted and directly measured stature. A previous study reported that Neolithic individuals were shorter than predicted from PRS, supporting the hypothesis of an environmental decrease in stature [11]. However, because they used PRS derived from European ancestry GWAS, their result can be confounded by stratification [26, 27]. Indeed, when they include ancestry in their regression models to control for the effects of population stratification, Neolithic individuals are no longer significantly shorter than expected. On the other hand, it is possible that including ancestry as a covariate, or even as a control for stratification at the GWAS level, could over-correct — removing real associations between SNP effects and ancestry. Results based on European GWAS are therefore difficult to interpret since they can lead to both false positive and false negative results.

To perform an unconfounded test of the Neolithic stature hypothesis, we used the East Asian derived PRS to compare differences in genetically predicted height across populations [31]. As well as being unconfounded, our analysis has a larger sample size than previous analyses (N=568 compared to N=167 in [11]). Following previous work[11], we estimate the environmental effects as the residuals from a regression of femur length on PRS and sex (Fig. 2A), although we note that these could also represent genetic effects not captured by the PRS. We divide our sample into pre-agriculture, agricultural Neolithic, and post-Neolithic time periods. We have defined these categories based on archaeological assessment of subsistence strategy rather than date or genetic ancestry (Methods), though we recognize that no classification system can fully categorize the complexity of subsistence strategies and transitions across the entirety of Europe.

**Figure 2:**
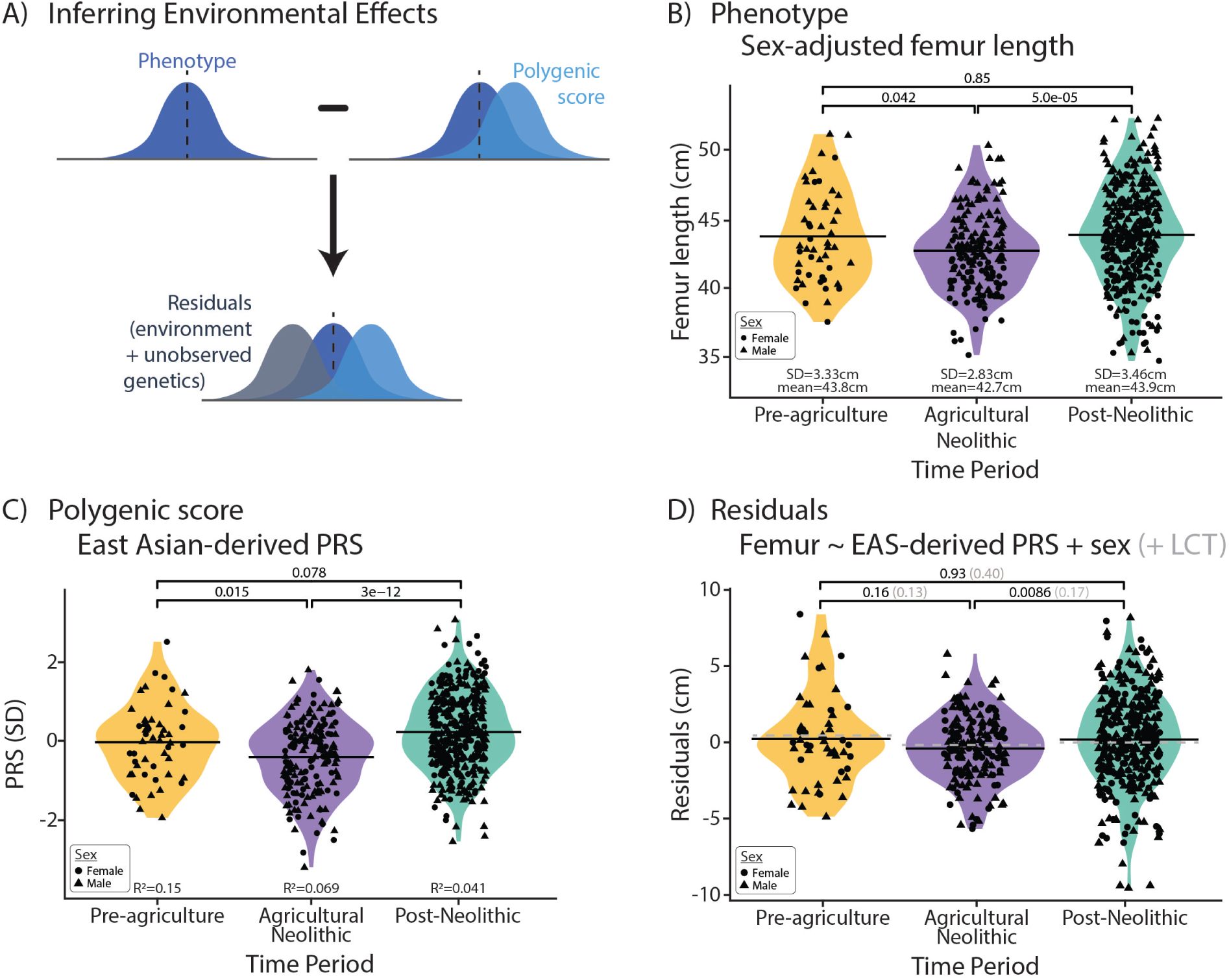
Were Neolithic individuals shorter than expected? A) Schematic illustration of inferring environmental effects. A phenotype is a combination of genetic and environmental effects. When we have known values for two of the three variables (phenotype and polygenic scores), we can reverse the equation to solve for the unknown quantity (environment). This inferred environmental component will also include the contribution of any unknown genetic effects. B) Phenotype effect: Femora are nominally shorter in the Neolithic than the pre-Neolithic and significantly shorter than the post-Neolithic period. Female femur lengths have been adjusted to remove the effect of sex by regressing out this variable; C) Polygenic score effect: PRS show that Neolithic populations are genetically shorter than the pre- and post-Neolithic; D)Residual effect: residuals from the linear regression of sex and East Asian-derived PRS indicate that compared to pre-Agriculture, Neolithic populations are not shorter than would be expected after accounting for these variables. Grey values and dotted lines indicate the effects of also including lactase persistence in the regression. P-values on all plots generated by two-tailed t-test.

We find that pre-agricultural femora were significantly longer than in the agricultural Neolithic (P=0.042, Fig. 2B). However, the absolute difference is small (0.9cm), corresponding to a difference of about 2.5cm in standing height. The polygenic score is significantly lower in the Neolithic (P=0.015, *β*=0.38SD, Fig. 2C), but the residuals are not significantly lower (P=0.16, Fig. 2D), in contrast to previous results [11]. While we do confirm a small decrease in average stature in the Neolithic, it appears to be at least partially genetic, limiting the potential for environmental effects.

Post-Neolithic populations are also slightly but significantly taller than Neolithic populations (1.1 cm in femur length, 3cm in stature, P=5.0 × 10*^−^*^5^, Fig. 2B). However, in this case, both the PRS and the residuals are significantly higher (by 0.63SD P=3.0 × 10*^−^*^12^ and 0.68cm P=0.009, respectively). The contrast in PRS indicates a substantial genetic contribution to the greater stature of post-Neolithic populations. Although higher residuals could reflect an environmental contribution, it could also represent the effect of genetic variation not captured by the PRS. Indeed, if we include lactase persistence in the regression (see below), the difference in residuals is no longer significant (P=0.17, Fig. 2D).

In summary, we cautiously refute the Neolithic stature hypothesis. Our data support that agricultural Neolithic populations were slightly shorter than both pre-agricultural and post-Neolithic populations but we find evidence for a genetic contribution to this difference, and limited evidence for any residual environmental effect. However, regardless of their origin, these average differences are small—2.5-3.5cm difference in stature compared to present-day differences of up to 7cm between European countries [32], global changes of up to 20cm in the past century [18] and differences of over 5cm between females from different Neolithic European populations [17].

### 2.3 Ancient association tests replicate modern GWAS effects

We next measured the effect of individual SNPs on stature in the ancient data. This dataset is too small to run a GWAS, so we focused on estimating effects at known height associated SNPs. Restricting to the 1000 most significant independent SNPs in the largest present-day height GWAS [22], we tested for association with femur length in our data. We reject a model where all 1000 effects are zero (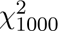 =1171, P=0.0001) and do not reject a model where the effect sizes in our data are identical to those in the present-day data (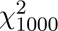 =1008, P=0.42). In fact, the chi-squared statistic for the second test is very close to the expected value (1000), and the slope from a regression of ancient on modern effect sizes is close to 1 (Fig. 3), suggesting that genetic effects on stature detected in present-day GWAS are very similar in ancient individuals. Of the 1000 SNPs, we replicate 81 in the correct direction at nominal significance (P=0.05) (Fig. 3). Consistent with the chi-squared test, this is significantly more than we would expect to replicate if we had no power (binomial test P=3.133 × 10*^−^*^5^) but is not significantly different than the 67 we would expect to replicate given the ancient sample size (binomial test P-value=0.081, Methods).

**Figure 3:**
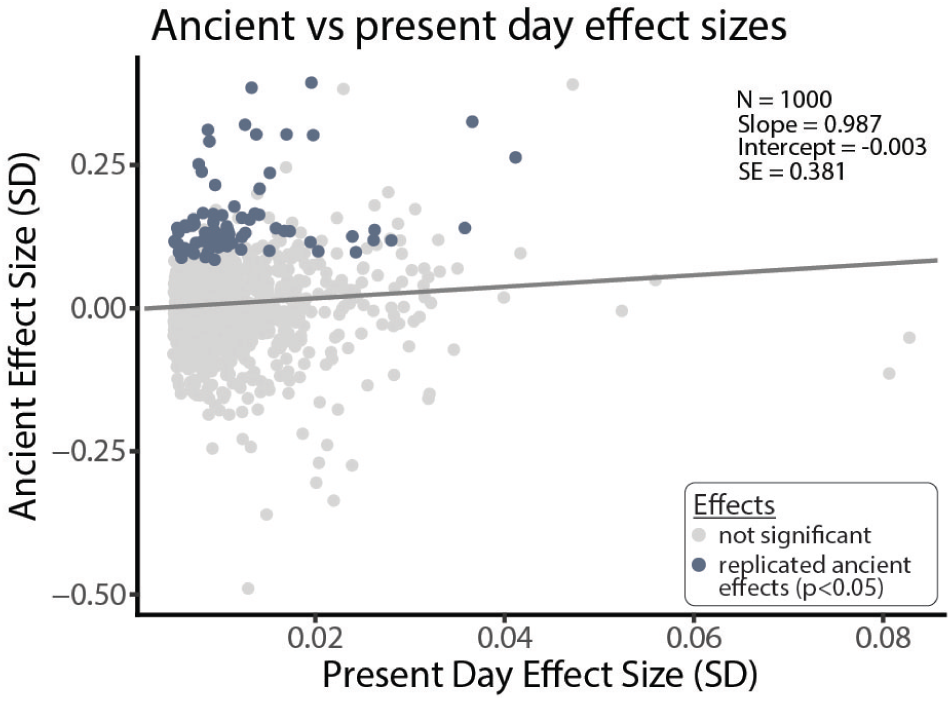
Re-estimating Ancient SNP effects. The relationship between present-day and re-estimated ancient effect sizes. Each point is one of the 1000 SNPs we tested. Grey points are not significant in our reestimation. Dark points are significant at P<0.05 with consistent direction. Regression line is based on Deming regression of modern ∼ ancient betas which takes into account the uncertainty in both dependent and independent variables

### 2.4 Lactase persistence had a large effect on ancient stature

Although SNPs detected in present-day GWAS have similar effects in ancient individuals, we hypothesized that the lactase persistence SNP rs4988235 could be associated with stature in ancient individuals even though it is not associated today [33]. Lactase persistence was under exceptionally strong selection between about 4500 and 1000 years BP [34], and must therefore have had a large phenotypic effect. The selective advantage remains unclear [35, 33], though it could be due to additional calories gained from digesting lactose [36, 37, 38, 39], or improved bone development via vitamin D and calcium metabolism [40, 41, 42]. After imputation about 20% of our ancient sample has either one or two copies of the lactase persistence allele (Fig. 4A). We find that the allele is significantly associated with longer femur length (P=6.01 × 10*^−^*^7^, *β*=0.358sd, se=0.071; Fig. 4A,B), even controlling for ancestry (P=2.38 × 10*^−^*^5^, *β*=0.318sd, se=0.075), and polygenic scores (P=3.02×10*^−^*^4^, *β*=0.256sd, se=0.071). As an alternative method of controlling for ancestry, we also analyzed just the post-Neolithic samples to reduce confounding due to the introduction of Steppe ancestry, with consistent results (P=0.0001, *β*=0.30SD, se=0.078; Fig. 4B) Confidence intervals for unimputed data overlap with those from the imputed genomes, indicating that this result is not due to imputation bias (Fig. 4B). Including lactase persistence in a regression of femur length on sex, ancestry and polygenic scores increases *R*^2^ by about 2% for both European and East Asian derived PRS, and removes the residual difference between Neolithic and post-Neolithic populations (Fig. 2D).

**Figure 4:**
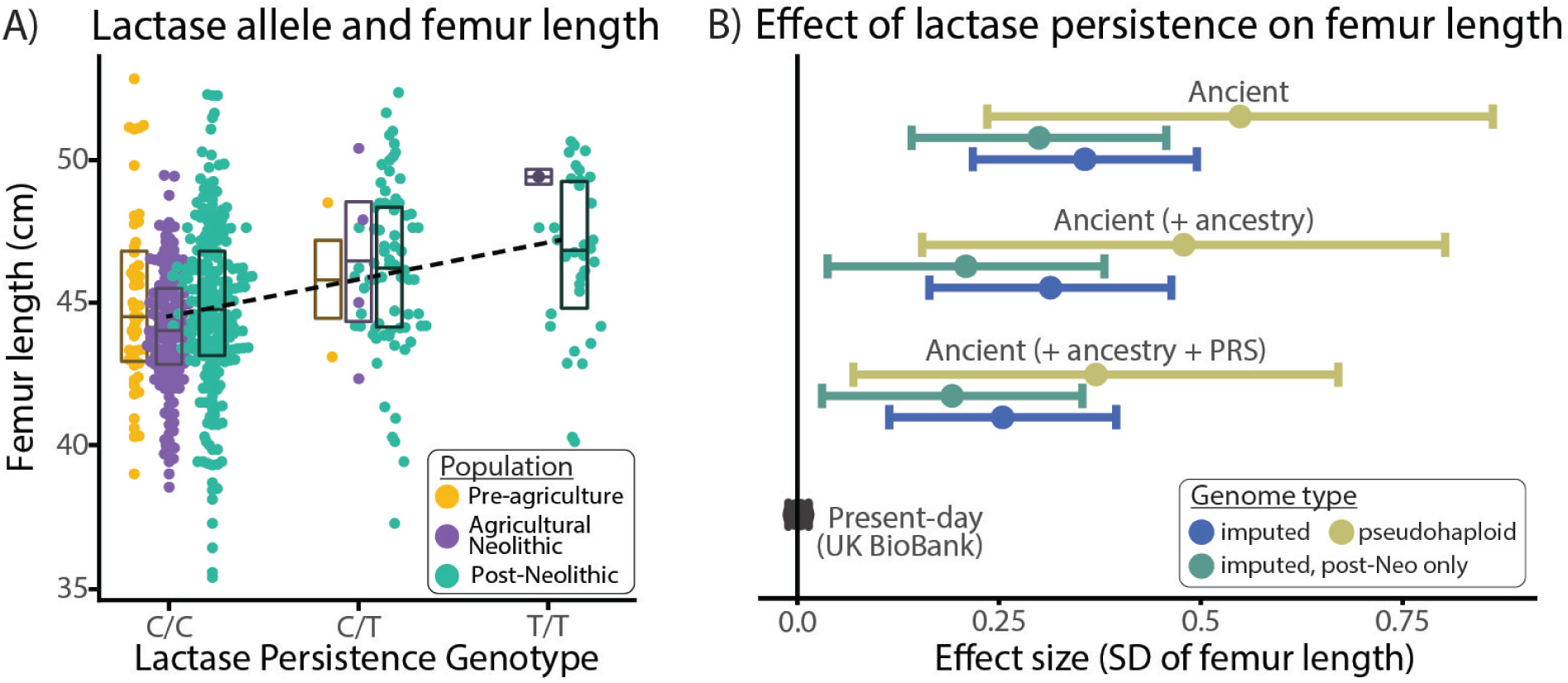
Effects of Lactase Persistence. A) Effect of lactase persistence on femur length. In present day GWAS there is no association between having the lactase persistence allele and being taller (top row). In our ancient samples, having the lactase persistence allele is significantly associated with heaving longer femora (second row; imputed: P=7.05×10^−8^, β=0.420cm/SD, se=0.077) with some ancestry-related variation (third row; imputed: P=3.43 × 10^−6^, β=0.372cm/SD, se=0.077). The effect is attenuated when we also include PRS in the model (calculated without chromosome 2 to avoid LD with the LCT gene) but is still significant (bottom row; imputed: P=0.0017, β=0.243cm/SD, se=0.079). Bars show standard error. B) Relationship between femur length and the lactase persistence (LP) allele. Having the LP allele significantly increases individual height (P=7.05 × 10^−8^) and presence of this allele is predominant in the post-Neolithic period. Boxes for each time period show quartiles and mean of femur length during those periods for each genotype. Dashed line is the regression line of femur length on LP allele. Femur lengths on the y-axis have been sex-adjusted by regressing out the effects of this variable.

## 3 Discussion

Our results are consistent with previous findings that the variance explained by PRS for height in ancient populations is about one fourth of the variance explained in present-day data [19, 11]. This likely reflects some combination of lower data quality (both genetic and phenotypic data), imputation accuracy, decline in performance due to genetic distance from the training population [28], and population stratification [43]. Population stratification is a particular concern when interpreting differences in polygenic score across populations, and for this reason we restrict our comparisons to PRS computed using East Asian GWAS summary statistics in which any stratification should be independent of European population structure.

Our results do not support a substantial decrease in stature caused by the European transition to agriculture [1]. First, though Neolithic populations were shorter than both Mesolithic and Bronze Age populations, these differences were small in both absolute and relative terms, especially when compared to variation among Neolithic [17] and modern [18] populations. Second, we find evidence that there was a significant genetic contribution to these changes, which means that the environmental effects, if any, were even smaller. Modern secular trends have shown that environmentally-driven shifts in stature can be dramatic, and these Neolithic changes were modest by comparison. It is therefore hard to support the idea that the adoption of agriculture had a substantial systematic effect on stature. On the other hand, the inconsistent results between studies [e.g., 6, 1, 4, 3, 2, 14] indicate that large, continent-wide surveys of Neolithic stature may be sensitive to sampling strategies and miss heterogeneous local patterns. Addressing these questions would then be better served by smaller regional analyses that can specifically focus on the complex interactions between hunter-gatherers, early farmers, and their cultural-environmental landscapes [17]. Finally, we note that although stature is generally regarded as a good proxy for population health [6, 44], there are other metrics of health (dental, paleopathology, demography) that we are not testing and it remains possible that the Neolithic transition was detrimental in ways that are not captured by stature.

We find that the effects of individual SNPs on stature are broadly the same in ancient populations as they are in the present-day. However, the lactase persistence associated SNP rs4988235 is an extreme exception. Although it has no effect on height (or any other substantive phenotype) today, we estimate its effect in ancient populations to be at least 0.25 standard deviations, corresponding to about 1.5cm in stature. This is an exceptionally large effect for a common variant, around 25 times larger than the most significant variants identified in present-day GWAS. If we were to run a GWAS for stature in Bronze or Iron Age Northern Europe, rs4988235 would likely be by far the most significantly associated SNP.

This has important implications for understanding the evolution of lactase persistence, one of the most intensively studied examples of recent human adaptation. Although the lactase persistence allele has been one of the most strongly selected variants in the entire human genome, there is still considerable debate about the nature of selection. It has recently been argued [33] that the lack of phenotypic associations with lactase persistence in the UK Biobank suggests the selective advantage was unrelated to the nutritional benefits of milk consumption. Our results illustrate the limitations of this argument— the absence of present-day effects cannot be used to reject prehistoric selective advantages. Instead, our findings provide support for lactase persistence having a nutritional advantage due to increased calories from digesting the lactose itself [36, 37, 38, 39] or improved vitamin D and calcium metabolism [40, 42].

In summary, our analysis demonstrates the value of integrating genomic and phenotypic data from ancient individuals. We hope that future ancient DNA studies will be more systematic about collating archaeological and anthropological data in order to enable more fine-scale analysis of phenotypic trends.

## 4 Materials and Methods

### 4.1 Genetic data

We used the genetic data from 568 ancient Eurasian individuals [45, 46, 47, 48, 49, 50, 51, 52, 53, 54, 55, 56, 57, 58, 59, 60, 61, 62, 9, 63, 64, 65, 10, 66, 67, 68, 69, 70, 71, 72, 12, 73, 74, 75, 76, 77, 78, 79, 80, 81, 34, 82, 83, 84, 85, 86, 87, 88, 89, 90, 91, 92, 93, 94, 95, 96, 97]. Most samples are genotyped by in-solution capture of a set of 1.24 million single nucleotide polymorphisms (SNPs), colloquially known as the “1240k” [9, 98]. For each individual, a single SNP was randomly selected at each of the 1240k sites, generating pseudohaploid data. For each sample, we downloaded the aligned bam files from the sources cited in their publications. For the few samples that did not have aligned bam files we aligned them using the pipeline from [50]. Calls were generated from aligned bams using the default parameters in *pileupCaller* (https://github.com/stschiff/sequenceTools). As these ancient samples are low coverage (median 0.51×), it is not generally possible to generate diploid genotypes. This potentially limits the performance of the polygenic scores, so we imputed diploid genotypes in our samples with *GLIMPSE* [20] using the ancient DNA pipeline developed by [21] and the 1000 Genomes Europeans [99] as reference panel.

There are a number of large height GWAS available from which we can test our ancient PRS predictions. We tested the height prediction performance of six different GWAS [23, 25, 24, 22] on both our imputed and unimputed samples. For each GWAS, we intersected the GWAS SNPs with those on the 1240k array, with the exception of those from the large [22] meta-GWAS. Due to data restrictions in the [22] analysis, they only published PGS weights for HapMap3 SNPs [100] from their full sample (N=approx 5 million), instead of traditional summary statistics. The intersection between HapMap3 and the 1240k array is low, and once the GWAS SNPs were clumped to calculate the PRS we did not have enough power to test it using our unimputed 1240k data. As an alternative, we tested PRS predictions from this GWAS by imputing the missing HapMap3 SNPs. Additionally, Yengo et al. also released traditional summary statistics from smaller GWAS, with less power, but fewer restrictions. These were successfully intersected with the 1240k array and clumped for testing both our imputed 1240k and unimputed data. In total, we tested PRS constructed from European GWAS from the UK Biobank [23] (n=7,008 SNPs), East Asian GWAS [24](n=3,326 SNPs), within-family sibling GWAS [25](n=1,115 SNPs), 1240k intersected Yengo Europeans (n=21,616 SNPs), 1240k intersected Yengo East Asians (n=1,940 SNPs), and 1240k intersected Yengo multi-ancestry (n=22,435 SNPs) SNP sets on both our imputed and unimputed 1240k ancient DNA data, as well as our HapMap3 imputed ancient DNA on PGS weights from the Yengo European (n=1,099,006 SNPs), East Asian (n=990,793 SNPs) and multi-ancestry (n=1,103,043 SNPs) meta-GWAS SNPs. We constructed PRS as previously described [19] using clumping and thresholding methods in *plink* v.1.9 (*--score*, p-value cut-off=10*^−^*^6^, *r*^2^=0.3, and 250kb windows). We chose parameters to maximize prediction based on [19] except for the larger Yengo meta-GWAS, from which the PRS were constructed without clumping, using the PGS weights published by those authors. For PRS calculations in the unimputed data, missing sites for each individual were ignored, effectively shrinking scores towards the sample mean.

We conducted principal component analysis (PCA) using *smartpca* [101] to project ancient individuals in to the PC space of 777 present-day Eurasian individuals [10]. Values from PCs 1-4 for each individual were included in regression analyses to control for ancestry related effects and are referred to collectively throughout the paper as “ancestry”. We included the date in years before present (BP) for each sample based on what was reported in the original publication for the sample. Whenever possible, we used ^14^C dates, otherwise we took the mid-point of the archaeological time range, and included genetic sex as reported in the original publications.

### 4.2 Osteological data

We obtained osteological metric data from publications as well as from direct measurements taken by the authors [102, 46, 103, 48, 104, 19, 105, 106, 107, 108, 109, 110, 111, 112, 17, 113, 114, 115, 116, 117, 118, 119, 120, 121, 122, 123, 124, 125, 126, 127, 128, 59, 129, 130, 131, 62, 132, 133, 134, 135, 136, 137, 138, 139, 140, 141, 142, 66, 143, 144, 145, 5, 68, 146, 11, 147, 148, 149, 150, 151, 152, 153, 80, 154, 155, 156, 157, 158, 159, 160, 161, 162, 88, 163, 92, 95, 164, 165, 166, 167, 168, 169, 170, 97, 171, 172, 173]. When only estimated statures have been published, rather than direct measurements, we reversed the equation of the cited stature estimation method to retrieve the maximum femur length that is associated with that stature [19]. There are a number of individuals (n=107) for which the estimation method used to calculate the published stature was not cited in the publication. In order to include these samples, we chose to estimate the femur length using the Trotter and Gleser (1952) equation as this is one of the most common methods, especially in publications predating [174].

We took dates for each sample from the mean of the calibrated ^14^C range in the original publication or the midpoint of the archaeological date range if no carbon date was available. Individuals were assigned to pre-agricultural, agricultural Neolithic, or post-Neolithic time periods based on archaeological assessments in publications associated with each sample. The pre-agricultural group includes all populations who do not yet have archaeological indications of agriculture, including hunter-gatherers with ceramics in Scandinavia. The post-Neolithic group is defined by cultures with metal technology and includes early Copper Age settlements. These lifestyle categories were not defined by genetic ancestry, although they are highly correlated (Fig. 1B).

### 4.3 Statistical models

**Evaluating PRS performance**. We fit linear models of femur length on PRS, sex, date in years BP, date-squared, and PCs 1-4 (ancestry) as described in [19]. For each linear model, we calculated the contribution of the PRS by taking the *R*^2^ of the full model with all variables and subtracting the *R*^2^ of a reduced model without the PRS variable; we refer to this simply as “*R*^2^” throughout the text [19]. For the analysis of femur-length as shown in Fig. 2B, we regressed out the effects of sex on femur length in order to remove the substantial sex effect, adjusting females to have the same mean as males. All p-values in Fig.2 come from two-tailed pair-wise t-test between time period groups.

**Re-estimating SNP effects**. We re-estimated the effects of the top 1000 [22] European GWAS SNPS by regressing each of the SNPs on femur length, including sex as a covariate (femur length SNP + sex). All SNPs have both the effect and other alleles present within the dataset. All missing 1240k SNPs were imputed as summarized above. Power for the re-estimation of PRS SNPs was calculated as

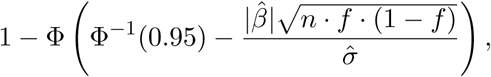

where Φ is the standard cumulative normal distribution function, *β* and *σ* are the estimated effect size and standard error of the UK Biobank GWAS, n is the ancient sample size (1136 haplotypes), and f is the minor allele frequency in the ancient sample. Because the UK Biobank GWAS is for height, not femur length, we adjusted beta and sigma to units of femur length by scaling them using the slope of the [174] stature estimation equation (2.7).

**Evaluating the effect of the lactase persistence allele**. To assess the effect of lactase persistence, we regressed femur length on sex in all models and also included genetic ancestry (PCs 1-4) and PRS where indicated. Given the number of SNPs involved in the height PRS, there is a strong likelihood that some of these could be in LD with the lactase gene (LCT) on chromosome 2, confounding the analysis. To mitigate this effect, we removed all SNPs on chromosome 2 from the GWAS summary statistics and re-calculated the PRS. For Fig. 4B we again adjusted female femur length to have the same mean as males.

### 4.4 Code and data Availability

All non-genetic data and polygenic scores used in this analysis are provided in Supplementary Table 1. Original ancient DNA data files can be downloaded from the resources provided in their cited publications and from the Allen Ancient DNA Resource (AADR). Previously published osteological data can be found in their cited sources which include the LiVES database (doi: 10.17171/2-12-2-1) and Dr. Christopher Ruff’s public dataset (https://www.hopkinsmedicine.org/fae/CBR.html.) R code for each analysis is available at https://github.com/mathilab/***.git.

## Acknowledgements

This work was supported by a grant from the National Science Foundation [BCS2123627] to IM. The content is the responsibility of the authors and does not necessarily represent the official view of the NSF or other funders.

